# Synergistic Antibiotic Activity and Integrated Genomic–Metabolomic Profiling of Patch - and Plaque-Derived *Staphylococcus aureus* in Mycosis Fungoides

**DOI:** 10.64898/2026.03.30.715345

**Authors:** Frederic Aurelius Straub, Laura Kim Serbin, Ilham El Barkani, Christiane Grünewald, Volker Mailänder, Nazzareno Dominelli

## Abstract

Mycosis fungoides (MF), the most common form of cutaneous T-cell lymphoma, is frequently accompanied by skin dysbiosis, with advanced lesions often dominated by multidrug-resistant *Staphylococcus aureus*. Increased *S. aureus* colonization is associated with clinical complications and accelerated disease progression, emphasizing the urgent need for effective antimicrobial strategies and a deeper understanding of bacterial adaptation to MF lesions.

Here, we evaluated synergistic antibiotic combinations and performed integrated phenotypic, genomic, and metabolomic profiling of MF-associated *S. aureus* isolates derived from patch and plaque lesions to understand virulence and pathogenicity driving mechanisms in microbe-host interactions. Several antibiotic combinations, most notably carbenicillin with either gentamicin or levofloxacin, exhibited strong synergy and restored antimicrobial activity against highly resistant strains. Comparative genomic analyses revealed that plaque-derived isolates carried expanded resistomes and virulence repertoires, including increased enterotoxins, immune-evasion, and stress-response factors, whereas patch-derived isolates encoded more genes linked to interbacterial competition, such as accessory components of the T7SS. Metabolomic profiles further supported these findings: plaque isolates produced metabolites linked to host interaction, dysbiosis, and inflammation, whereas patch isolates showed profiles consistent with ecological competition.

In summary, this work provides insight into the distinct adaptation strategies of *S. aureus* across MF disease stages. The differential virulence and resistance repertoires observed between patch- and plaque-derived isolates suggests progressive adaptation toward the host microenvironment, potentially influencing disease progression and patient outcomes. Additionally, our findings identify synergistic antibiotic combinations as promising therapeutic approaches for targeting multiresistant MF-associated *S. aureus*.

**Importance Statement:** The role of multidrug-resistant *Staphylococcus aureus* in worsening clinical outcomes of mycosis fungoides remains poorly understood, despite its frequent dominance in advanced lesions. Bridging the gap between clinical observation and microbiological mechanisms is essential for clarifying how *S. aureus* (SA) persists within MF skin and for identifying therapeutic alternatives for SA-positive patients, where treatment options remain limited. This study sheds light on two major clinical needs: the lack of effective antibiotic strategies and the limited insight into bacterial factors that may accelerate MF progression. By integrating synergistic antibiotic testing with genomic, phenotypic, and metabolomic profiling, our work provides insight into stage-specific adaptation patterns of MF-associated *S. aureus*. These findings identify promising therapeutic directions and establish a framework for future studies to understand the role of *S. aureus* in MF pathogenesis and exploring how its effects may be therapeutically mitigated.

## Introduction

Mycosis Fungoides (MF) is a non-Hodgkin cutaneous T-cell lymphoma (CTCL) that primarily affects the skin and lymphatic system (1). This disease accounts for 50-70% of all CTCL cases (2). In Europe, the prevalence is estimated to be 2.6 cases per 10,000 people, predominantly affecting adults aged 50-60 years and is more prevalent in men (3). MF is clinically categorized into stages IA to IVB based on disease progression (4). Early MF presents with patches, defined as irregularly reddened, itchy skin areas (5), and a subset of patients progresses to the more advanced stages - plaques, and tumours (6–8). In the final stage, MF may evolve into Sézary syndrome with high numbers of malignant T cells in the blood (9, 10). Currently, treatment is stage dependent, as in early stages, a variety of skin directed therapies (SDTs) are employed, while in more advanced stages, systemic therapies are utilized (11–13). These therapies often provide only short-term benefits, and long-term responses remain uncertain (12). MF significantly impacts skin health, particularly the cutaneous microbiome. Patients exhibiting persistent cutaneous inflammation and T-cell dysregulation often display a more disrupted skin microbiome (14, 15). Studies found that patients with a high abundance of *Staphylococcus* experience increased erythroderma and pain, which may be linked to T helper cell polarization in colonized areas due to *staphylococcal* enterotoxins. This polarization potentially drives inflammation and dysregulation, thereby exacerbating disease progression (14, 16).

Additionally, MF-patients often exhibit a high abundance of *S. aureus* on their skin or mucosal areas (17). High *S. aureus* levels stratify the skin microbiome and may accelerate MF progression (18). Furthermore, increased *S. aureus* colonization is associated with lymphoma promotion, likely due to its enterotoxin production (16, 19).

*S. aureus* is a Gram-positive, facultative anaerobic and opportunistic pathogenic bacterium (20). As a member of the ESKAPE group of pathogens, it has gained notoriety for its fast acquisition of multidrug resistance (21, 22). Specifically, *S. aureus* strains resistant to methicillin (MRSA), or vancomycin (VRSA) represent a major challenge in clinical environments and are among the leading causes of nosocomial infections (23). Upon infection, *S. aureus* can cause a wide spectrum of diseases, including endocarditis, bacteremia, osteomyelitis, and pneumonia, many of which are associated with high mortality rates (23–26). Accordingly, it is considered one of the most significant contributors to antimicrobial resistance and infection-related mortality on a global scale (27, 28). Moreover, its pathogenicity is enhanced by a broad range of virulence factors, such as biofilm formations, toxins, surface proteins and immune evasion strategies (29, 30).

This study aimed to elucidate the genomic architecture, resistance profiles, and metabolic features of patch- and plaque-derived *S. aureus* isolates to uncover virulence factors linked to MF progression and dysbiosis. In parallel, we assessed synergistic antibiotic combinations as potential therapeutic options against multiresistant plaque-associated *S. aureus*.

## Results

### Isolation of multiresistant *S. aureus* strains from MF-patients’ skin plaques

Skin swabs from multiple MF plaque sites yielded several *Staphylococcus* species, with *S. aureus* being isolated as the dominant organism. Given the clinical relevance of *S. aureus* in MF, representative isolates from lesional (*S. aureus* MFMZ4-1, MFMZ4-3) and non-lesional (*S. aureus* MFMZ4-2) skin, together with the previously described plaque isolate MFMZ1 (18), were subjected to antimicrobial susceptibility testing and were prioritized for in-depth analyses. Several strains exhibited a multidrug-resistant phenotype. All isolates showed resistance to β-lactam antibiotics, while lesional strains exhibited broader resistance profiles than the non-lesional isolate. The plaque-derived isolate MFMZ1 showed intermediate susceptibility to erythromycin, gentamicin, levofloxacin and SXT, while plaque isolates *S. aureus* MFMZ4-1 and MFMZ4-3 exhibited additional resistance to gentamicin and levofloxacin, with MFMZ4-3 also resistant to erythromycin and tetracycline.

In contrast, non-lesional isolate *S. aureus* MFMZ4-2 remained largely susceptible, except for intermediate susceptibility to levofloxacin. Importantly, reference strain *S. aureus* DSM11823 displayed rifampicin resistance, whereas all MF-derived isolates remained susceptible to this antibiotic. All strains retained susceptibility to last-line antibiotics tigecycline and ceftarolin (Tab. 1).

**Table 1:** Antibiotic resistant profiles of *S. aureus* plaque patient’s isolates MFMZ1, MFMZ4-1, MFMZ4-2 and MFMZ4-3 and control strain DSM11823. Susceptibility of *S. aureus* isolates were investigated via plate diffusion assay and evaluated according to EUCAST standard. *S. aureus* DSM11823 strain served as control. R = resistant, I = intermediate, S = sensitive/susceptible. Three biological replicates were performed.

| Tested antibiotic | *S. aureus* MFMZ1 | *S. aureus* MFMZ4-1 | *S. aureus* MFMZ4-2 | *S. aureus* MFMZ4-3 | *S. aureus* DSM11823 |
| --- | --- | --- | --- | --- | --- |
| Ampicillin 25 µg | R | R | R | R | R |
| Carbenicillin 10 µg | R | R | R | R | R |
| Ceftarolin 5 µg | S | S | S | S | S |
| Erythromycin 15 µg | I-S | S | S | R | S |
| Gentamicin 10 µg | I-S | R | S | R | I-R |
| Levofloxacin 5 µg | I | R | I | R | I |
| Linezolid 30 µg | S | S | S | S | S |
| Penicillin G 2U | R | R | R | R | R |
| Rifampicin 5 µg | S | S | S | S | R |
| SXT (23.75 µg + 1.25 µg)<br>*Sulfamethoxazole + Trimethoprim* | I | S | S | I | S |
| Tetracycline 30 µg | S | S | S | R | S |
| Tigecycline 15 µg | S | S | S | S | S |

### MIC assay reveals synergistic effect of antibiotics combination against multiresistant *S. aureus* isolates

Minimum inhibitory concentrations (MICs) of the selected antibiotics were determined for the *S. aureus* isolates MFMZ1, MFMZ4-1, MFMZ4-2, MFMZ4-3 and the reference strain DSM11823. Ampicillin showed no inhibitory activity against any isolate. For *S. aureus* MFMZ1, MICs of 100 µg/ml for gentamicin (Gent) and 150 µg/ml for kanamycin (Kan) were determined, while carbenicillin (Carb) inhibited growth only at 2000 µg/ml. *S. aureus* MFMZ4-1, MFMZ4-2 and control strain DSM11823 showed similar susceptibility profiles, with gentamicin being most effective (MIC 50 µg/ml), followed by kanamycin (75 µg/ml), whereas carbenicillin required ≥ 250 µg/ml for growth inhibition. In contrast, *S. aureus* MFMZ4-3 was inhibited only by gentamicin (200 µg/ml), with no detectable inhibition by kanamycin or carbenicillin within the tested range. Methicillin MICs were 1.56 µg/ml for *S. aureus* MFMZ4-1, MFMZ4-2 and DSM11823, and 3.125 µg/ml for MFMZ1. No methicillin-mediated inhibition of MFMZ4-3 strain was observed up to 200 µg/ml (Fig. 1).

**Figure 1:**
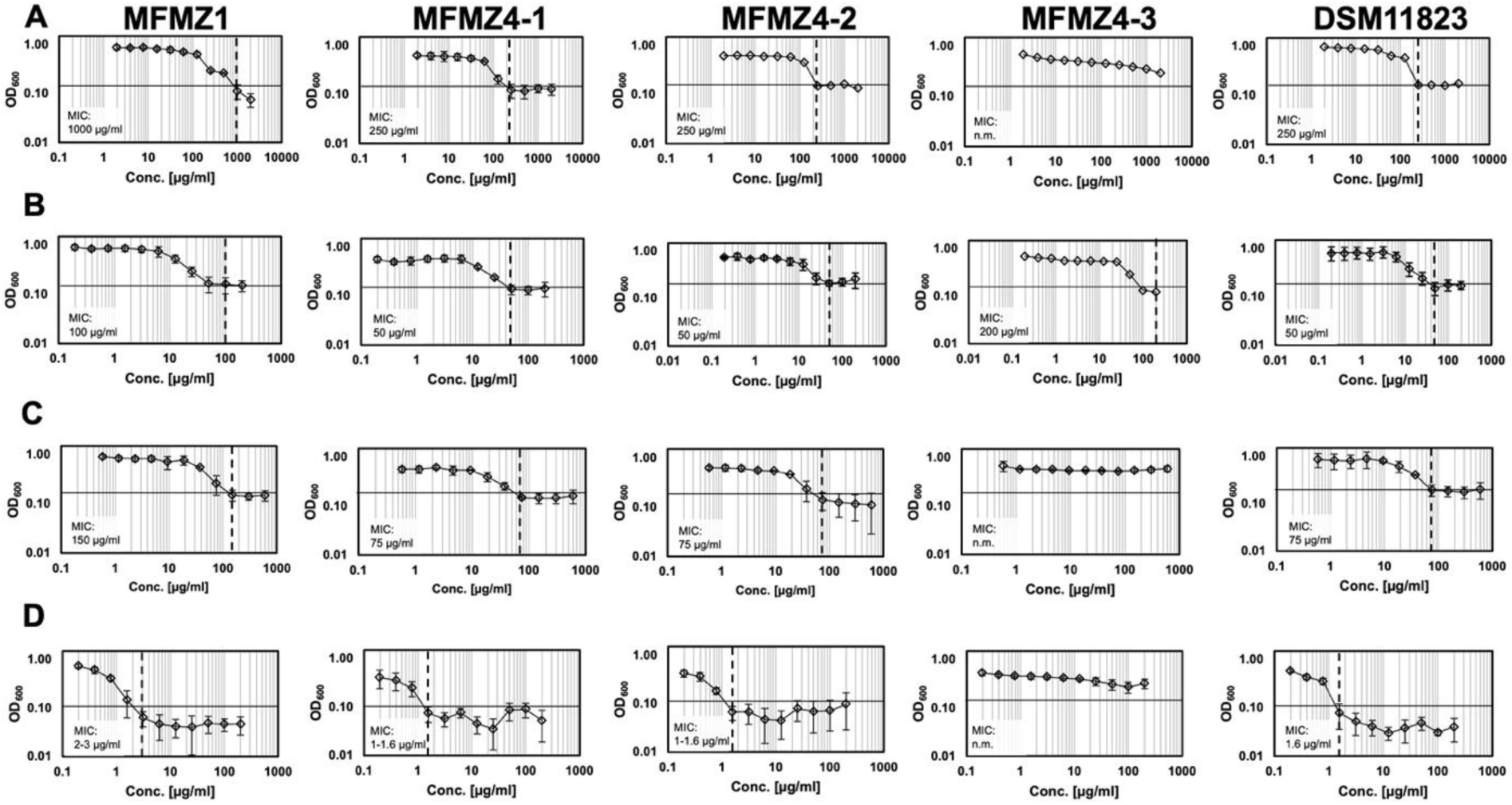
Minimal inhibitory concentration (MIC) profiles of β-lactam and aminoglycoside antibiotics against MF-associated *S. aureus* isolates. (A) Carbenicillin, (B) gentamicin, (C) kanamycin, and (D) methicillin were tested in 96-well plates across serial concentrations. From left to right, isolates MFMZ1, MFMZ4-1, MFMZ4-2, MFMZ4-3, and the control strain DSM11823 are shown. Growth was monitored by OD₆₀₀ measurements. The horizontal black line indicates the initial medium OD₆₀₀ baseline, and the vertical dashed line indicates the MIC. “n.m.” indicates that the MIC was not measurable. At least three biologically independent replicates were performed.

Further, biofilm formation was investigated to assess potential effects of antibiotics exposure, especially in sub-MIC concentrations. Biofilm formation remained low, except for carbenicillin-treated *S. aureus* MFMZ4-1 and MFMZ4-2, which showed increased biofilm production at concentrations near the MIC (Fig. S1). *S. aureus* MFMZ4-3 did not form detectable biofilm under any tested condition.

Synergy testing by combining β-lactams with aminoglycosides revealed no synergistic effect for ampicillin and kanamycin (FIC 0.5625 to >2.0). Likewise, the combinations Carb/Kan and Carb/Gent did not exhibit synergy against strain MFMZ4-3 (FIC > 2.0 and FIC > 1.25, respectively). In contrast, all other antibiotic combinations demonstrated additive or synergistic effects. Notably, although Amp alone exhibited no inhibitory effects, the Amp/Gent combination displayed clear synergy against *S. aureus* MFMZ4-3, markedly reducing MIC values (Amp 125 µg/ml + Gent 50 µg/ml; FIC < 0.5). The strongest synergistic inhibition was observed against *S. aureus* DSM11823, MFMZ4-1, and MFMZ4-2 (MICs 312.5 µg/ml Amp + 12.5 µg/ml Gent; FIC 0.3125), whereas MFMZ1 strain required 62.5 µg/ml Amp + 25 µg/ml Gent (FIC 0.375) for inhibition (Fig. 2). Except for *S. aureus* MFMZ4-3 the combination Carb/Kan displayed additive to synergistic activity in all isolates (FIC ∼0.5), with particularly strong synergy against *S. aureus* MFMZ1 (FIC 0.01). The Carb/Gent combination showed synergy against most strains with substantially reduced MICs and FIC values ranging from 0.13 to 0.5 (Fig. 2). The fluoroquinolone levofloxacin (Lev) was subsequently tested alone and in combination with Carb and/or Gent (Fig. 3). Lev exhibited inhibitory activity against all strains, with MIC values of 0.8 µg/ml for *S. aureus* MFMZ4-1, and MFMZ4-2, while DSM11823 strains required 1.6 µg/ml. In contrast, *S. aureus* MFMZ1 required a higher concentration (8 µg/ml), while MFMZ4-3 demonstrated markedly reduced susceptibility, with a MIC of 50 µg/ml (Fig. 3). Combining Lev with Carb, Gent, or both reduced the required inhibitory concentrations across all strains. Synergistic activity of Lev/Carb was observed exclusively against all strains, with FIC indices ranging from 0.3 to 0.45, except for *S. aureus* DSM11823 (FIC 0.55, additive effect). The Lev/Gent combination exhibited synergy only against *S. aureus* MFMZ1 (FIC 0.23) accompanied by substantial reductions in MIC values, while solely an additive effect against *S. aureus* MFMZ4-3 and DSM11823 strain could be observed (Fig. 4).

**Figure 2:**
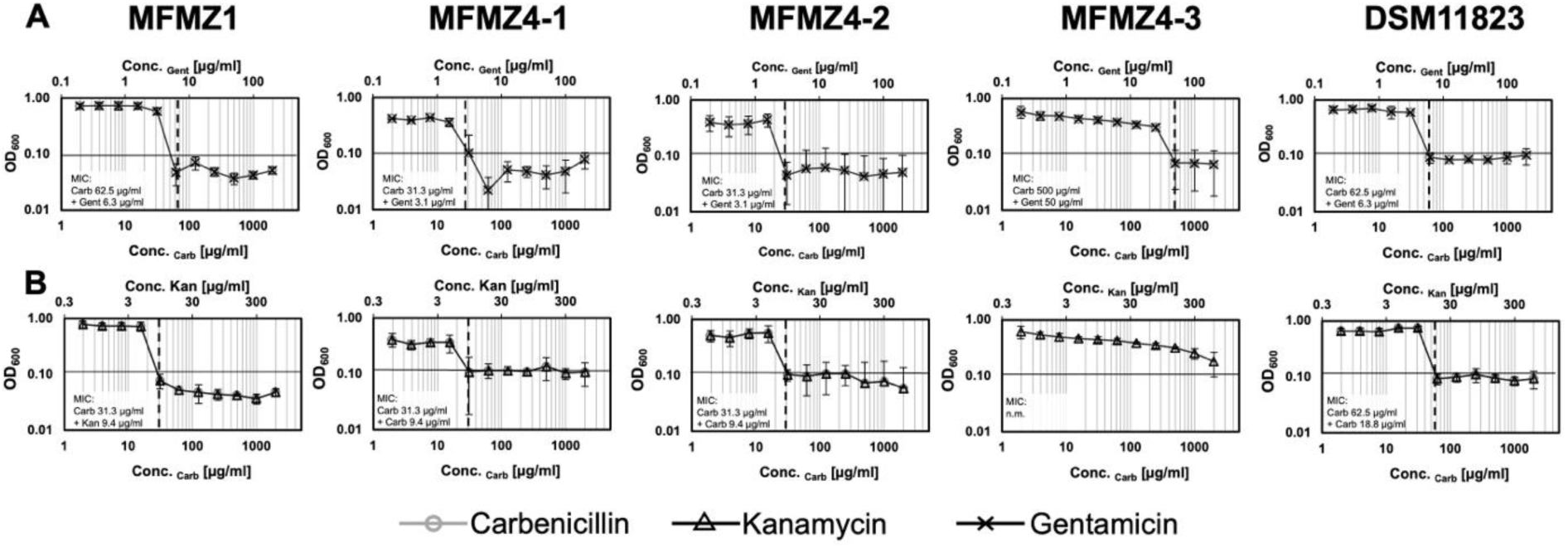
Synergy and minimal inhibitory concentration (MIC) profiles of β-lactam and aminoglycoside antibiotic combinations tested against MF-associated *S. aureus* isolates. (A) Combinatory effect of carbenicillin (grey line with circle) with gentamicin (black line with X), and (B) combinatory effects of carbenicillin with kanamycin (black line with triangle). From left to right, isolates MFMZ1, MFMZ4-1, MFMZ4-2, MFMZ4-3, and the control strain DSM11823 are shown. Growth was monitored by OD_600_ measurements. The horizontal black line indicates the initial medium OD₆₀₀ baseline, and the vertical dashed line indicates the MIC. “n.m.” indicates that the MIC was not measurable. At least three biologically independent replicates were performed.

**Figure 3:**
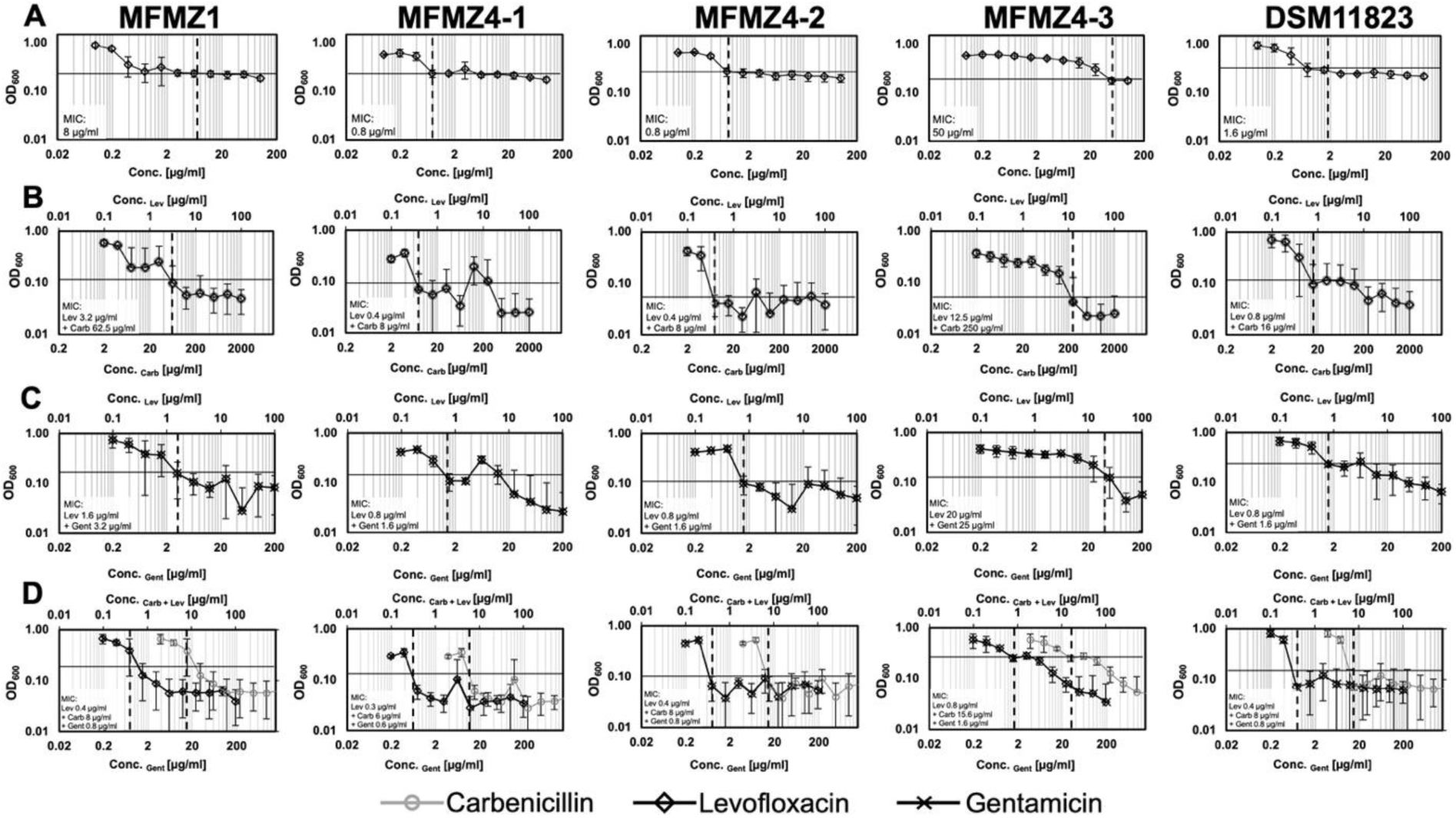
Minimal inhibitory concentration (MIC) profile of fluoroquinolone levofloxacin and synergy of combinations with β-lactam and aminoglycoside tested against MF-associated *S. aureus* isolates. (A) Levofloxacin (black line with diamond) alone, and in combination with (B) carbenicillin (grey line with circle), (C) gentamicin (black line with X), and (D) with carbenicillin and gentamicin. From left to right, isolates MFMZ1, MFMZ4-1, MFMZ4-2, MFMZ4-3, and the control strain DSM11823 are shown. Growth was monitored by OD_600_ measurements. The horizontal black line indicates the initial medium OD_600_ baseline, and the vertical dashed line indicates the MIC. At least three biologically independent replicates were performed. At least three biologically independent replicates were performed.

**Figure 4:**
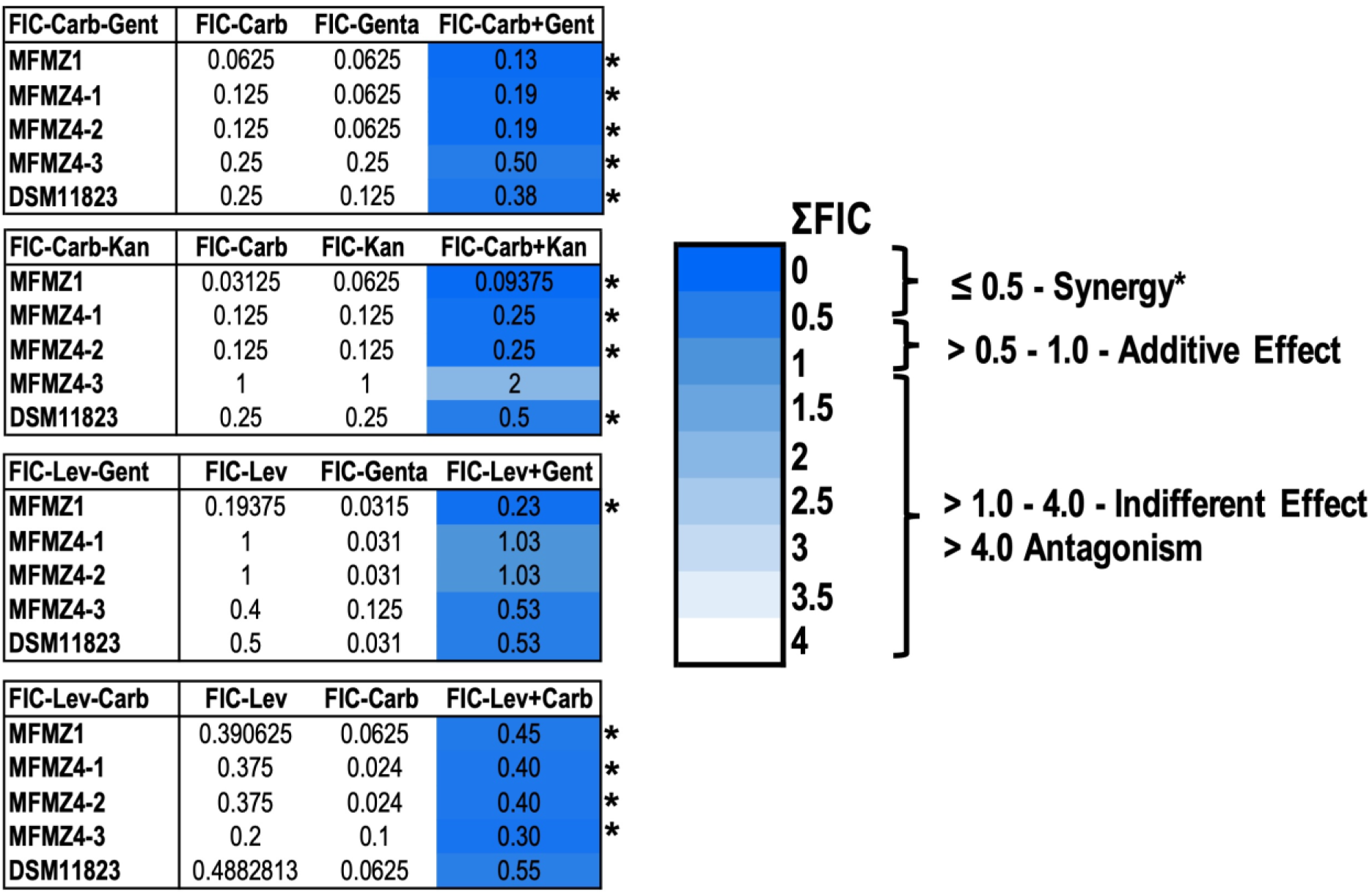
Fractional inhibitory concentration (FIC) indices of antibiotic combinations against MF-associated *S. aureus* isolates. For top to bottom FIC values for carbenicillin (Carb) – gentamicin (Gent), carbenicillin (Carb) – kanamycin (Kan), levofloxacin (Lev) – gentamicin (Gent), and levofloxacin (Lev) – carbenicillin (Carb) combinations are shown for *S. aureus* isolates MFMZ1, MFMZ2-1NL, MFMZ2-2L, MFMZ4-1, MFMZ4-2, MFMZ4-3, and control strain DSM11823. Tables display the individual FIC values for each antibiotic and the combined FIC (ΣFIC). The heatmap indicates the ΣFIC classification as defined on the right panel: ≤ 0.5 indicates synergy, > 0.5–1.0 additive effects, > 1.0–4.0 indifferent effects, and > 4.0 antagonism.‘*‘ indicates synergy. At least three biologically independent replicates were performed.

### Genomic Comparison Reveals Enhanced Virulence and Resistance Repertoires in Plaque-associated *S. aureus*

Comparative genomic analysis revealed marked differences between patch-derived and plaque-derived *S. aureus* isolates (Tab. 2; Fig. 5). Patch isolates MFMZ2-1NL and MFMZ2-2L shared a largely conserved virulence gene repertoire, including four exotoxin clusters, one exfoliative toxin cluster, two stress-response islands (LexA and a putative inosine-5′-monophosphate dehydrogenase) and an intact type VII secretion system (T7SS) locus. *S. aureus* MFMZ2-2L additionally harboured a membrane-bound metalloendopeptidase island absent from non-lesional isolate. In contrast, plaque patient derived isolates (MFMZ1, MFMZ4-1, MFMZ4-2, MFMZ4-3) exhibited greater genomic heterogeneity, with a broader repertoire of pathogenicity islands. All harbour conserved stress-adaptation systems (GroES/EL operon and Clp protease clusters), multiple exotoxin- and superantigen-islands, the Excalibur Island (CrcB_1) and one T7SS locus. The MFMZ4 isolates additionally carried lactococcin-related gene-islands and the TIGR-TAS bacterial immunity system within the T7SS locus (Fig. 5A). Functional presence/absence profiling further highlighted these differences. Patch isolates lacked multiple virulence factor genes, like toxins/hemolysins (e.g. *hlgBC*), adhesins (*sdrD*, *clfB*), immune-evasion (*sak*, *scn*), and the Spl protease operon whereas plaque isolates displayed largely complete virulence gene sets across categories (Fig. 5A–C). A notable difference between isolates was observed for genes associated with T7SS (Fig. 5B). Both patch-derived strains (MFMZ2-1NL and MFMZ2-2L) and plaque-derived MFMZ1 encoded a comparatively high number of T7SS-related genes, especially multiple *ess* and *esx* homologs. In contrast, MFMZ4 isolates carried markedly fewer T7SS-associated loci. Distribution of enterotoxin and superantigen genes also differed markedly (Fig. 5C): Patch-patient derived strains lacked several classical enterotoxin genes (*sec*, *sei, sem*, *sep*), present in plaque-patient derived strains, some of which present in higher number (Fig. 5C). All strains, especially patch-patient derived strains harbour an extensive set of superantigen-like proteins. Additional virulence determinants only found in plaque-derived include, among others, hyaluronate lyase HysA, staphylokinase Sak, *staphylcococcal* complement inhibitor SCIN, and the surface anchored adhesin SdrD, the latter only occurred in truncated form in non-lesional *S. aureus* MFMZ2-1NL. All isolates harbour the clumping factor coding *clfA*, whereas *clfB* was absent from MFMZ2-1NL. Protein A (Spa), also present in all strains, displays an expanded octapeptide repeat region, especially in plaque isolates.

**Figure 5.**
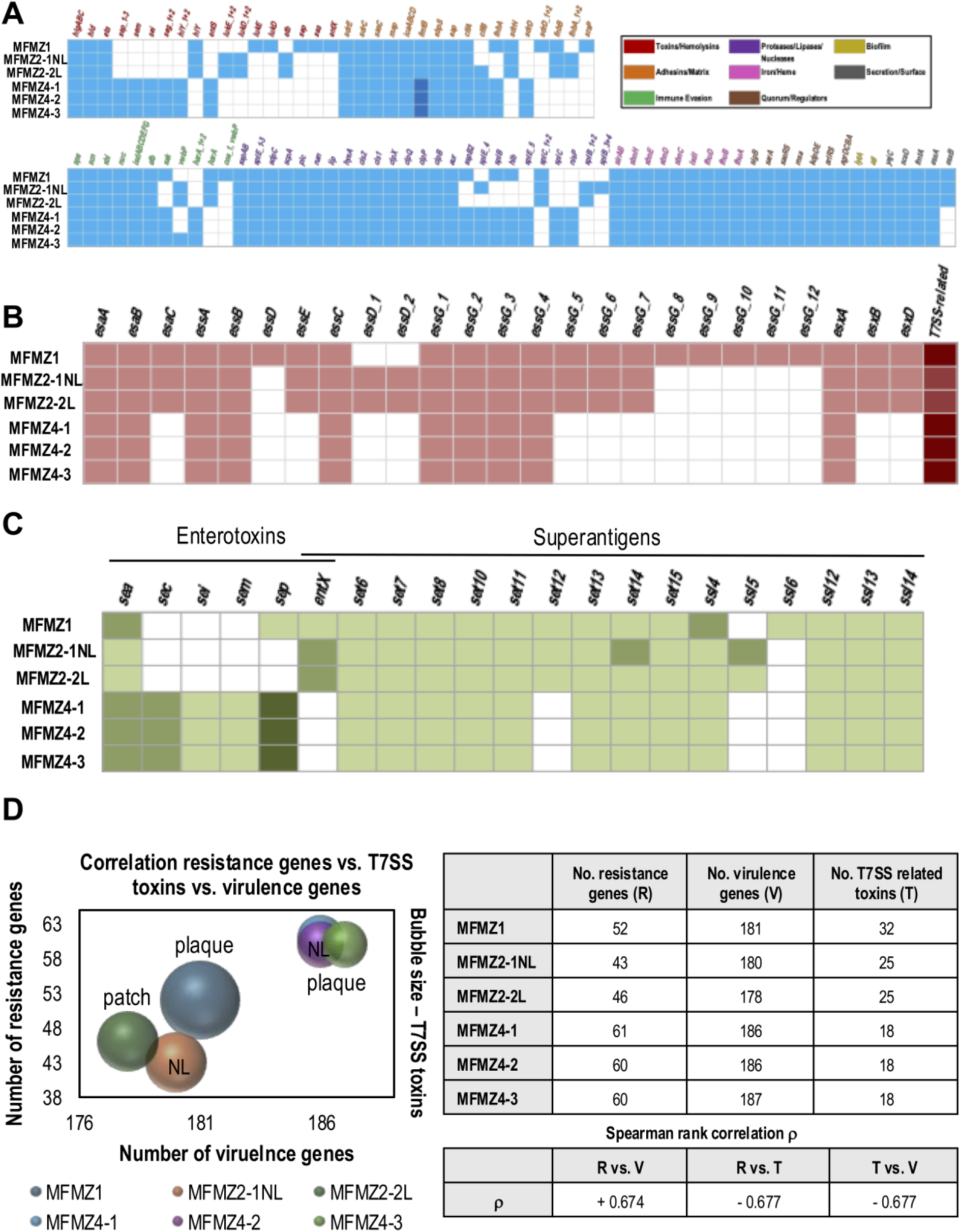
Comparative genomic and virulence-associated features of MF-associated *S. aureus* isolates. Presence–absence heatmap overview of (A) pathogenicity-associated genomic islands across isolates MFMZ1, MFMZ2-1NL, MFMZ2-2L, MFMZ4-1, MFMZ4-2, and MFMZ4-3. Font colours of genes indicate functional categories as defined in the legend on the top right panel. (B) T7SS related virulence-associated genes, and (C) distribution of enterotoxins and superantigen-like genes. Overall, darker shades indicate a higher number of genes present, no colour indicates absence. (D) Correlation analysis of resistance genes (R), virulence factors (V), and T7SS-related toxins (T). Left panel: bubble plot illustrating the number of resistance genes versus virulence genes, with bubble size representing the number of T7SS-associated toxins. Right panel: table summarizing gene counts and Spearman correlation coefficients.

**Table 2:** Overview of virulence-associated genomic features in MF-associated *S. aureus* isolates. Virulence factors and genomic islands were identified from whole-genome sequencing data from *S. aureus* isolates MFMZ1, MFMZ2-1NL, MFMZ2-2L, MFMZ4-1, MFMZ4-2, and MFMZ4-3 using NCBI, PHASTER, and IslandViewer. Numbers in brackets indicate the number of the respective factor identified; „Isl.“ denotes islands.

| Virulence factor / Cluster | MFMZ1 | MFMZ2-1NL | MFMZ2-2L | MFMZ4-1 | MFMZ4-2 | MFMZ4-3 |
| --- | --- | --- | --- | --- | --- | --- |
| **Genomic islands** |  |  |  |  |  |  |
| co-chaperone (GroES/EL) & Clp protease; leukocidin/hemolysin | Island 1 | - | - | Island 2 | Island 4 | Island 10 |
| Enterotoxins (superfamily) | - | - | - | Island 3 | Island 3 | Island 9 |
| Excalibur island (extracellular Ca-binding, NUDIX, CrcB_1) | Island 3 | - | - | Island 4 | Island 1 | Island 8 |
| Lactococcin (bio-synthesis & transport) | - | - | - | Island 11/13 | Island 5/6 | Island 1/2 |
| LexA-associated island + other genes (e.g., SCP/PR1-like) | - | Island 10 | Island 1 | - | - | - |
| Membrane-bound metallo-endopeptidase & truncated MHC II analog protein | - | - | Island 7 | - | - | - |
| Pathogenicity island proteins (misc.) | - | - | - | Island 9 | Island 8 | Island 4 |
| Phage-related proteins / loci | - | Isl. 4/8/9 | Isl. 6/3 | Island 6 | Island 10 | Island 6 |
| Putative IMP dehydrogenase, PemK, YjdF | - | Island 2 | Island 10 | - | - | - |
| Bacteriocin immunity protein (hemin-regulated ABC transporter) | Island 3 | - | - | Island 1 | Island 1 | Island 1 |
| Superantigens | - | Isl. 1/2/6/7/8 | Isl. 4/10 | Island 3 | Island 3 | Island 3 |
| T7SS loci | (1) | (1) | (1) | (1) Island 10 | (1) Island 7 | (1) Island 3 |
| Multidrug ABC transporter / TRAP (RNAIII activating protein) | Island 2 | - | - | Island 4 | Island 2 | Island 8 |
| **Outside of genomic islands** |  |  |  |  |  |  |
| Exotoxin clusters | (6) | (4) | (4) | (3) | (4) | (4) |
| Exfoliative toxins | - | (1) | (1) | - | - | - |
| Prophages | (3) | (3) | (4) | (2) | (2) | (2) |
| Superantigens | (16) | (12) | (12) | (12) | (18) | (16) |
| TIGR-TAS system (bacterial immunity) | - | - | - | (1) | (1) | (1) |

Analysis of antimicrobial resistance (AMR) genes revealed substantial differences between isolates (Tab. S2, Fig. 5D). The plaque derived strain *S. aureus* MFMZ1 strain harboured 52 AMR-associated genes, including multiple determinants for β-lactam resistance (*bla* genes, *fmtC*, *abcA*), lincosamide resistance (*lnrL*), glycopeptide resistance (*tca* and *vra*), fluoroquinolone resistance (*norABC*), among others. The MFMZ4 plaque isolates harboured even larger AMR repertoires (60 - 61 genes) comprising additional macrolide resistance determinants (*emrB*, *qacA*), aminoglycoside resistance genes (*msbA*, *aac*(3)), and further β-lactam–associated loci. Notably, *S. aureus* MFMZ4-3 carried an additional copy of *norB*, resulting in the highest number of NorB-type efflux components across all isolates. In contrast, patch-patient derived isolates encoded fewer AMR genes: 43 genes in non-lesional MFMZ2-1NL and 46 in lesional MFMZ2-2L isolates, while still retaining the core multidrug-resistance gene repertoire (Table S2). Across all isolates, the total number of AMR genes positively correlated with the overall virulence genes (Spearman ρ = 0.67), while the abundance of T7SS-associated virulence genes showed no positive association with either AMR gene counts or total virulence factors, and displaying a negative correlation (ρ = –0.67 for both comparisons) (Fig. 5D).

Lastly, antiSMASH analysis identified secondary metabolite biosynthetic gene clusters (SMBs) in all *S. aureus* isolates. Patch-patient derived strains harbour six SMBs, while seven SMBs were found in the remaining isolates. All strains shared a conserved type III polyketide synthase (T3PKS) region of unknown function. Five additional SMBs associated with aureusimines production, the metallophores staphylopine, and staphyloferrin A/B, and the Agr quorum sensing system were identified. The additional detected SMB in plaque-derived strains differed among the isolates: *S. aureus* MFMZ1 harboured a listeriolysin synthesis-associated region, whereas MFMZ4 strains encoded a class II lanthipeptide gene cluster.

### GC-MS analysis reveals distinct metabolic signatures in MF-associated *S. aureus* isolates

GC–MS profiling revealed distinct metabolite patterns across the isolates (Fig. 6). Phosphonic acid was detected in all strains except *S. aureus* MFMZ4-3. The MFMZ4-1 isolate uniquely produced the polyamine putrescine, whereas *S*.

**Figure 6:**
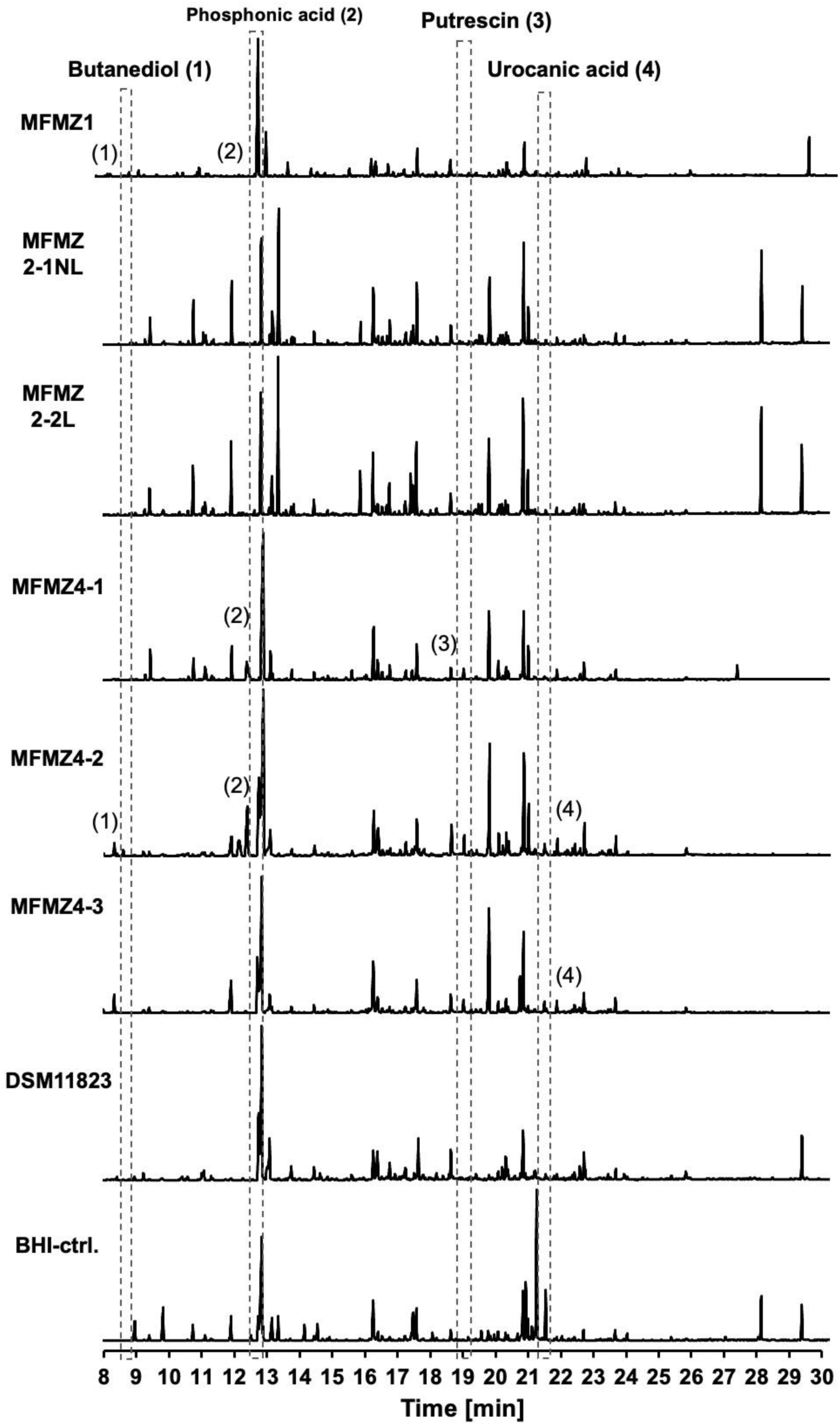
GC–MS chromatograms of supernatants from MF-associated *S. aureus* isolates. From top to bottom chromatograms of *S*. *aureus* isolates MFMZ1, MFMZ2-1NL, MFMZ2-2L, MFMZ4-1, MFMZ4-2, MFMZ4-3, control strain DSM11823, and the medium control (BHI-ctrl) are shown. Dashed vertical lines mark the retention time ranges of identified metabolites: (1) 2,3-butanediol, (2) phosphonic acid, (3) putrescine, and (4) urocanic acid. Numbers correspond to annotated peaks in the chromatograms. Time is shown on the x-axis in minutes. At least three independent biological replicates were performed.

*aureus* MFMZ1 and MFMZ4-2 produced measurable amounts of 2,3-butanediol. In addition, urocanic acid was identified in the supernatants of *S. aureus* MFMZ4-2 and MFMZ4-3. Overall, the GC–MS analysis demonstrated clear strain-specific metabolic signatures among the MF-associated isolates.

In summary, our combined phenotypic and genomic analyses highlight pronounced differences between patch- and plaque-derived *S. aureus* isolates, reflected in their resistome, genomic virulence, biosynthetic gene clusters, and metabolite production. In addition, we could demonstrate several synergistic antibiotic combinations with therapeutic potential against multi-resistant *S. aureus*. Together, these findings provide insights into strain-specific adaptations of MF-associated *S. aureus* that may contribute to disease progression.

## Discussion

MF-patients with advanced disease progression frequently exhibit a dysbiotic skin microbiome dominated by multiresistant *S. aureus* (18). In this study, we demonstrate that several antibiotic combinations show robust synergistic activity against multiresistant MF-associated *S. aureus*. We further provide an integrated phenotypic, genomic, and metabolic characterization of isolates derived from patch and plaque lesions. Together our findings indicate determinants for strain-specific adaptations that may facilitate persistence and progression withing the MF skin environment.

By combining different antibiotics from different classes, we could determine synergy and could restore or markedly enhance antimicrobial activity, against isolates with high-level resistance to mono-use. Particularly, β-lactam combinations with aminoglycosides or fluoroquinolones, such as carbenicillin with gentamicin or levofloxacin, displayed strong synergistic effects and substantially lowered MIC values, including in highly resistant isolates like *S. aureus* MFMZ4-3. This observation is in line with previous studies reporting that antimicrobial activity could be either restored or enhanced, when β-lactams are paired with aminoglycosides or fluoroquinolones, where mono-application was insufficient, therefore resulting in synergistic effects (31, 32). Notably, combinations such as carbenicillin with gentamicin or levofloxacin substantially reduced MICs in several isolates, including *S. aureus* MFMZ4-3, for which MICs were barely detected. These findings support the idea that the multi-target inhibition reduces the effectiveness of individual resistance mechanisms, which might also increase the genetic barrier to resistance (33).

Synergistic effects involving fluroquinolones are of particular interest given the increasing fluoroquinolone resistance (34), as restoring their activity would be therapeutical valuable due to its reported anti-biofilm properties (35–37). Although, overall biofilm formation was rather low, carbenicillin exposure near the MIC increased biomass in *S. aureus* MFMZ4-1 and MFMZ4-2, which is in accordance with the observation that sub-inhibitory β-lactam levels may induce biofilm formation and persister emergence (38, 39). These observations highlight the importance of dosing strategies that prevent sub-MIC exposure to prevent survival or formation of a tolerant subpopulation and treatment failure (40, 41).

Further genomic comparisons revealed pronounced differences between patch- and plaque-derived isolates. Plaque isolates carried expanded AMR repertoires, including β-lactam resistance genes, lincosamide and glycopeptide resistance determinants, and multiple fluoroquinolone efflux systems, classifying all strains as multidrug-resistant (42). This corresponds to the reduced susceptibility and elevated MICs across multiple antibiotic classes. Beyond AMR, plaque isolates exhibited broader sets of pathogenicity islands (PIs) like GroES/GroEL which facilitate refolding of misfolded proteins, and Clp proteases which contribute to proteostasis, antibiotic stress adaptation, and virulence regulation (43–45). This might support tolerance to antimicrobials such as tetracyclines (43), and stabilize resistance proteins like efflux pumps (44, 46). GroES/GroEL is further employed by *S. aureus* in stress adaption against heat or acid stress (47). This may support bacterial survival under the conditions given by the fluctuating skin microenvironment of MF-patients, as functional redundancy among stress adaption mechanisms can provide survival advantages under extreme conditions or when one mechanism is compromised (48). Increased PI diversity as found in plaque-associated strains suggests an improved colonization capacity by enhancing adaptability to host-specific niches (49, 50). Interestingly, T7SS, typically associated with interbacterial competition (51, 52), differed between isolate groups. While all strains encoded intact T7SS loci, T7SS-associated genes such as *essG* were reduced in plaque-derived isolates, whereas patch isolates and MFMZ1 strain carried more extensive T7SS effector sets. Enlarged T7SS loci, have been linked to enhanced bacteria- and host-targeting functions with increased virulence (53, 54). The loss of specific T7SS-effectors EssG in plaque isolates coincided with enrichment of TIGR-TAS (Tandem Interspaced Guide RNA-associated proteins) immunity genes within their T7SS loci, a system associated with mobile element interference, interbacterial competition, or genetic inheritance, and may also complement CRISPR-Cas immunity (55, 56). This pattern suggests a putative ecological shift: in early-stage patch lesions, with higher microbiome diversity, *S. aureus* may rely on competitive mechanisms, whereas plaque-associated strains appear more adapted to long-term host interaction (57).

Additional virulence factors, including HysA, Sak, SCIN, or Spa were detected across all strains, and may contribute to MF-associated colonization. HysA hyaluronidase facilitates tissue invasion by degrading hyaluronic acid (58, 59). Staphylokinase (Sak) with fibrinolytic and proteolytic activity, and staphylococcal complement inhibitor SCIN, co-located in immune evasion clusters (60, 61), protect the pathogens against phagocytosis also during early biofilm formation (62, 63). Plaque isolates displayed expanded Spa repeat regions, which may modulate IgG binding and host–pathogen interactions (64, 65).

Analysis of biosynthetic gene clusters (BGCs) provided further evidence for strain-specific adaptation. All isolates harboured conserved clusters for aureusimines and siderophores (staphyloferrins A/B), which promote host interaction and iron acquisition in nutrient-limited environments (66, 67). Additional BGCs in plaque isolates, such as class II lanthipeptide clusters, may further contribute to niche adaptation and interbacterial dynamics. Similar lantibiotics have been linked to bactericidal activity (68). Further, investigating *agr* QS cluster revealed predominance of *agr* type III, although in other skin inflammatory diseases such as atopic dermatitis (AD), *agr* type I is most prevalent (69). This variability may reflect ecological selection, as distinct *agr* systems can inhibit each other, as studies showed that *S. aureus* (*agr I*) QS was inhibited by *S. hominis* harbouring *agr* type III (69–72). However, distribution patterns may also vary by geographic region (73, 74).

Metabolite profiling further supported these genomic trends. All identified compounds implicate stress- and biofilm-associated shifts across the isolates. Phosphonic acid, present in all isolates except *S. aureus* MFMZ4-3, serves as an alternative phosphor source, that may indicate metabolic flexibility under nutrient or antibiotic stress (75). Putrescine in *S. aureus* MFMZ4-1 is associated with biofilm formation, membrane stability, and the response to acid stress (76). Production of 2,3-butanediol in MFMZ1 and MFMZ4-2 strains suggests roles in redox homeostasis and putative stress resistance, antibiotic tolerance or biofilm formation (77). Both might contribute to an overall increased virulence and persistence of *S. aureus* using different strategies. Urocanic acid, which usually functions as a skin moisturizing factor with antibacterial properties, was produced by *S. aureus* MFMZ4-2 and MFMZ4-3 (78). In high concentrations urocanic acid might be harmful for the host, as it modulates immune responses, particularly when converted to immunosuppressive cis-urocanic acid upon UV exposure. It acts as an immunosuppressive molecule that dampens Langerhans cell activity and T-cell responses (79, 80), which may theoretically contribute to MF disease progression.

Together, these findings reveal distinct adaptation strategies among MF-associated *S. aureus* and support the therapeutic potential of synergistic antibiotic combinations. The divergence between patch- and plaque-derived isolates indicates a shift from interbacterial competition in early-stage lesions toward increased host-adaptation and virulence in advanced disease. Overall, our integrated phenotypic, genomic and metabolic analyses provide a framework for understanding and further studying how *S. aureus* may contribute to MF disease progression. Here we highlight several factors that might be involved in disease progression and evasion of immunity and therapeutics that must be further to improve clinical management of MF.

## Material and Method

### Isolation and identification of MF-associated *Staphylococcus aureus* strains

To isolate *S. aureus* from mycosis fungoides patients, sterile cotton swabs were used to sample the skin. Samples were collected from patients treated at the Department of Dermatology, University Medical Centre Mainz (Germany). Swabs were taken from lesional skin - patches and plaques - and from non-lesional areas, preferably the healthy counterpart of the respective lesion. For this study, swabs were collected from patches on the calf of patient 2024/01/18_001 and from plaques on the cheek of patient 2024/03/01_004. These samples were streaked onto blood agar plates [1% (w/v) peptone from casein, 1% (w/v) meat extract, 0.5% (w/v) NaCl, 50 ml sterile sheep blood] and onto *Staphylococcus-*selective Vogel-Johnson agar plates (Sigma Aldrich). After incubation at 37°C, single colonies were picked and subjected to species identification by colony PCR using *tuf*-specific primers [*tuf* fwd – GGCCGTGTTGAACGTGGTCAAATCA & *tuf* rev TIACCATTTCAGTACCTTCTGGTAA) (81)] and the OneTaq® polymerase (NEB) following the manufacturer’s protocol. Successful amplification was assessed via agarose gel electrophoresis using a 1% (w/v) agarose gel in TAE buffer [40 mM Tris/HCl, 20 mM glacial acetic acid, 1 mM EDTA] containing 0.2 mg/ml Midori Green (Nippon Genetics), run for 20 min at 110 V. Visible amplicons were excised and purified using the HiYield® PCR Clean-Up & Gel Extraction Kit (SLG Gauting) according to the manufacturers protocol. Sanger sequencing of the purified fragments was performed by StarSeq® GmbH (Mainz) using the respective primers. Resulting sequences were analysed with the Basic Local Alignment Search Tool BLAST® (https://blast.ncbi.nlm.nih.gov/Blast.cgi) to determine the bacterial species.

### Cultivation of *S. aureus* strains

To grow overnight cultures of *Staphylococcus aureus*, one single colony was inoculated into liquid tryptone-soy-broth (TSB) [1.7% (w/v) tryptone; 0.5% (w/v) NaCl, 0.3% (w/v) soy peptone tryptone, 0.3% (w/v) meat extract, 0.25% (w/v) glucose, 0.25% K_2_HPO_4_] and cultivated at 37°C under aerobic conditions. For agar plates 2% (w/v) agar was added to TSB media. For this study isolates from the patients were used as well as previously isolated *S. aureus* MFMZ1 strain (18).

### Antibiotic diffusion assay

To characterize the respective *S. aureus,* antibiotic sensitivities and resistances, overnight cultures were prepared and adjusted to an OD600 = 1.5. After that, 1 ml of the diluted culture were pipetted into 50 ml of already melted and hand warm 0.8% (w/v) TSB soft agar. This mixture was then directly poured into sterile petri dishes. After agar solidification, antibiotic disc dispensers (BD BBL^TM^ Sensi-Disc^TM^ and Oxoid™ Blank Antimicrobial Susceptibility discs) were used to distribute antimicrobial susceptibility test discs (Tab. S1). Those plates were firstly incubated at 4°C for 2 h to allow diffusion of the antibiotics through the agar and subsequently incubated at 37°C.

### Minimal Inhibitory Concentration (MIC) and antibiotic synergy

Minimum inhibitory concentrations (MICs) of the selected antibiotics were determined for the *S. aureus* isolates MFMZ1, MFMZ4-1, MFMZ4-2, and MFMZ4-3, as well as for the reference strain DSM11823, to provide the basis for subsequent synergy analyses.

Overnight cultures of the respective strains were prepared in brain-heart infusion (BHI) broth and adjusted to an OD_600_ of 0.1 in 2x media. Two-fold serial dilutions of respective tested antibiotics were prepared in 96-well plates with sterile water and bacteria were added, respectively. The concentration range tested for the antibiotics were 500 µg/ml - 0.977 µg/ml ampicillin, 2000 µg/ml – 1.953 µg/ml carbenicillin, 200 µg/ml – 0.195 µg/ml gentamicin, 600 µg/ml – 1.172 µg/ml kanamycin and 100 µg/ml – 0.098 µg/ml levofloxacin. All of them were tested alone and in combinations: β-lactam antibiotics ampicillin and carbenicillin with aminoglycosides gentamycin or kanamycin; and the fluoroquinolone levofloxacin with β-lactam carbenicillin and/or aminoglycoside gentamicin. Additionally, MIC for methicillin (200 µg/ml – 0.195 µg/ml) was determined. Absorbance was measured using Tecan Spark microplate reader at OD_600_ at timepoints 0 h and 24 h. The plates were incubated at 37°C under aerobic conditions. Three biological independent replicates were performed. Afterwards MICs were determined and resulting fractional inhibitory effects (FICs) were evaluated to determine potential synergy of the antibiotic combinations (82–84). Additionally, biofilm production was quantified (85–88). After incubation, the plates were tapped out to remove culture broth and planktonic cells followed by a washing step with water and dried for 30 min. Afterwards, the plates were stained with 135 µl 1% (v/v) crystal violet solution for 30 min. The plates were tapped out and washed with water to remove excessive crystal violet solution. After drying the plates overnight, 135 µl of 30% (v/v) acetic acid was added to the wells, to dissolve the crystal violet bound to the biofilm. Absorbance was measured using the microplate reader at 575 nm (86).

### Whole genome sequencing of MF-associated *Staphylococcus aureus* isolates

To examine genetic differences between *S. aureus* strains isolated from lesions and non-lesional areas from MF-patients, whole genome sequencing (WGS) was performed. Therefore, sample were prepared according to the instructions of MicrobesNG (Brimingham, UK). Briefly, a single colony of the respective strain was picked and resuspended in 200 µl sterile phosphate-buffered saline (PBS) [137 mM NaCl, 2.7 mM KCl, 10 mM Na_2_HPO_4_, 1.8 mM KH_2_PO_4_, pH 7.2]. Then, 100 µl were transferred to 25 ml TSB medium for liquid cultivation at 37°C and 100 µl were plated onto TSB agar plates. Once reaching the exponential phase at an OD_600_ of 0.75, the cells were harvested via centrifugation at 4,500 rpm for 15 min. The cell pellet was resuspended in 1 ml of inactivation buffer (provided by MicrobesNG) and sent for sequencing. WGS occurred using Illumina short reads (2x250 bp) with 60x target coverage and bioinformatically analysed. Sequencing data were provided as contigs, which were further investigated using multi-draft-based scaffolder 1.6 (MeDuSa) including 450 randomly chosen *S. aureus* reference genomes via NCBI dataset tool for assembly (89), following quality control and contamination removal of raw reads with Kraken 2 (90). Via further BLAST analyses un-assembled reads were identified as plasmids. Afterwards, Prokka was used to annotate the *S. aureus* genomes, identifying genes and functional elements using genus-specific databases and Rfam RNA data (91). For enhanced annotation accuracy, the genomes were analysed using an HMM-based protein annotation tool and UniProt accession mapping. Genome comparison was conducted with the *nucmer* program from the MUMmer 4.0 package (92). Furthermore, the genomes were examined with additional tools to identify further genetic factors: Antibiotic resistance genes were identified using Abricate with the CARD (93), ResFinder (94), and MEGARes (95) databases, as well as AMRFinderPlus (96) was used. Mobile genetic elements and genomic islands were detected using gnomAD (97), and IslandViewer 4 (98), respectively. Virulence factors were identified using the VFDB databases (99). For identification of secondary metabolite biosynthesis gene clusters antiSMASH was used (100). After completing the automated analyses, the annotation tables were manually reviewed to identify the respective genetic factors that were not detected by the applied tools. Sequence data have been deposited under the BioProject PRJNA1443228 and are associated with the following BioSamples: SAMN56718705 (*S. aureus* MFMZ1), SAMN56718706 (*S. aureus* MFMZ2-1NL), SAMN56718707 (*S. aureus* MFMZ2-2L), SAMN56718708 (*S. aureus* MFMZ4-1), SAMN56718709 (*S. aureus* MFMZ4-2), and SAMN56718710 (*S. aureus* MFMZ4-3).

### Gas Chromatography-Mass Spectrometry (GC-MS) Analysis

To characterize the secondary metabolite profiles and to further understand virulence of the respective MF-*S. aureus* isolates, 50 ml of brain-heart infusion (BHI) broth were inoculated with cells of each strain and incubated aerobically at 37 °C for 72 h. Cultures were centrifuged twice at 4,500 rpm for 30 min to remove bacterial cells. The resulting supernatants were sterile-filtered using a Steriflip® vacuum filtration system equipped with a Millipore Express® PLUS membrane (0.22 µm) and subsequently subjected to GC/MS analysis. For GC/MS analysis, samples were derivatized according to Koek et al., 2006 (101). Briefly, 50 µl of each sample was lyophilized, and derivatization was carried out by adding ethoxyamine hydrochloride and pyridine, followed incubation for 90 min at 40°C. Subsequently, samples were silylated for 50 min at 40°C with 70 µl MSTFA. GC/MS analysis was performed using a Shimadzu GCMS-QP2010 system equipped with a quadrupole mass spectrometer. 5 µl aliquots of each sample were injected in split mode (1:10) into a DB5-MS capillary column (30m x 250µm I.D., 0.25 µm film thickness, Phenomenex, Germany) using an AOC20i autosampler at an injection temperature of 250°C. The initial oven temperature was set to 70°C; five minutes after injection, the temperature was increased at a rate of 10 °C/min to 320°C and held for 10 min. Helium was used as the carrier gas at a flow rate of 1 ml/min. Detection was achieved using MS in electron-impact ionization mode with full-scan acquisition (15–800 m/z). The ion source temperature was set to 200°C and the transfer line to 250 °C. Compound identification was achieved by comparing the obtained mass spectra with entries in the NIST spectral library.

## Acknowledgment

We thank Ralf Heermann for fruitful scientific discussions. We also thank Alina Carolin Breunig and Janne Marie Sowa for their assistance with preparatory laboratory tasks.

